# Downregulation and delayed induction of photosynthesis by coordinated transcriptomic changes induced by sink-source imbalance

**DOI:** 10.1101/2023.01.19.524789

**Authors:** Yui Ozawa, Aiko Tanaka, Takamasa Suzuki, Daisuke Sugiura

**Author notes:** Corresponding author: Daisuke Sugiura. Other authors.

## Abstract

Understanding comprehensive mechanisms of the downregulation of photosynthesis induced by accumulation of non-structural carbohydrates (NSCs) is essential for the future food security.x Despite numerous studies, whether NSCs accumulation directly affects steady-state maximum photosynthesis and photosynthetic induction, as well as underlying gene expression profiles, remains unknown so far.

We evaluated the relationship between photosynthetic capacity and NSCs accumulation induced by cold-girdling, sucrose feeding, and low nitrogen treatment in *Glycine max* and *Phaseolus vulgaris*. In *G. max*, changes in transcriptome profiles were further investigated focusing on physiological processes of photosynthesis and NSCs accumulation.

NSCs accumulation decreased maximum photosynthetic capacity and delayed photosynthetic induction in both species. In *G. max*, such photosynthetic downregulation was explained by coordinated downregulation of photosynthetic genes involved in Calvin cycle, Rubisco activase, photochemical reactions, and stomatal opening. Furthermore, sink-source imbalance may have triggered a change in the balance of sugar-phosphate translocators in chloroplast membranes, which may have promoted starch accumulation in chloroplasts.

Our findings provided an overall picture of the photosynthetic downregulation and NSCs accumulation in *G. max*, demonstrating that the photosynthetic downregulation is triggered by NSCs accumulation and cannot be explained simply by N deficiency.

**One Sentence Summary:** Accumulation of nonstructural carbohydrates directly induced both downregulation and delayed induction of photosynthesis by coordinated transcriptomic changes in photosynthetic genes in *Glycine max*.

## Introduction

It has been pointed out that photosynthetic downregulation is triggered by accumulation of non-structural carbohydrates (NSCs) such as soluble sugars and starch in leaves, which would be more pronounced by elevated CO_2_ and low nitrogen availability (Sage et al., 1989; Drake et al., 1997). Since leaf photosynthesis is a key determinant of crop productivity, the downregulation of photosynthesis would have a negative impact on crop yields in the future. Therefore, a comprehensive understanding of the mechanisms of sugar-induced downregulation could contribute to solving the future food problems through improved crop productivity.

In general, NSCs accumulate in source leaves when the sink capacity is lower than the source capacity (Kasai, 2008; Sugiura et al., 2017). The transcript levels of photosynthetic genes such as *RbcS* encoding Rubisco small subunits and other key Calvin cycle enzymes were suppressed by the accumulation of NSCs, resulting in the decreased CO_2_ assimilation rate (Nie et al., 1995; Krapp and Stitt, 1995; Van Oosten and Besford, 1995; Moore et al., 1999). Other study reported that NSCs accumulation also causes the cell wall thickening, which contribute to the decrease in mesophyll conductance (Mizokami et al., 2019; Sugiura et al., 2020). The downregulation of photosynthesis is suggested to become more remarkable by the limited root growth due to small pot size (Arp, 1991) and the decreased sink activity due to low nitrogen (N) availability (Ruiz-Vera et al., 2017). On the other hand, it is suggested that the downregulation of photosynthesis could be explained simply by N deficiency rather than NSCs (Geiger et al., 1999). This is because elevated CO_2_ often cause nitrogen deficiency due to imbalance between CO_2_ assimilation and N uptake in plants, leading to the decreased contents of chlorophyll and photosynthetic proteins containing abundant N (Stitt and Krapp, 1999). Previous study showed that the Rubisco content is selectively decreased while maintaining leaf N content occurs in response to NSCs accumulation in *Phaseolus vulgaris* and *Chenopodium album* (Sage et al., 1989; Nakano et al., 1998; Sugiura et al., 2019), while it has been suggested that the decrease in the Rubisco content was simply due to N deficiency in *Triticum aestivum* and *Oryza sativa* (Nakano et al., 1997; Theobald et al., 1998). However, since causal relationship between NSCs accumulation and decreased leaf N content has not been clarified by these studies, it remains unknown whether NSCs directly causes the photosynthetic downregulation. Therefore, it is necessary to establish an experimental system that can evaluate the effect of NSCs and N deficiency separately.

Another issue is whether photosynthetic induction is also affected by the accumulation of NSCs remains unknown since most previous studies on the photosynthetic downregulation have focused on the steady-state photosynthesis. Photosynthetic rates rarely reach the steady-state maximum in natural environments with fluctuating light regimes (Pearcy, 1990; Naumburg and Ellsworth, 2002; Yamori, 2016), and the gradual increase in photosynthetic rate with increased light intensity is called photosynthetic induction (Walker, 1973). Photosynthetic induction is characterized by three steps: (1) activation of the ribulose-1,5-bisphosphate (RuBP) regeneration process, (2) activation of Rubisco, and (3) stomatal opening (Pearcy, 1990). The first phase is completed within 2 minutes of increased light intensity, and the cyclic electron flow around photosystem I and ion transporters/channels localized in the thylakoid membrane are involved (Yamori and Shikanai, 2016; Carraretto et al., 2016). The second phase will be completed within 5 to 10 minutes after the Rubisco activase is fully activated (Yamori et al., 2012; Carmo-Silva and Salvucci, 2013; Kaiser et al., 2016), while the third phase takes an hour to reach a steady state depending on the plant species (McAusland et al., 2016) since CO_2_ uptake by stomatal opening is the most limiting factor of photosynthesis. Although previous study reported increased crop biomass production and yield in the Free-air CO_2_ enrichment (FACE) experiments, it is pointed out that they were far less than expected from the growth chamber experiments (Ainsworth and Long, 2005; Leakey et al., 2009). This suggests that the NSCs accumulation delayed the photosynthetic induction responses under fluctuating environments in the FACE experiments compared with the growth chamber experiments with constant light environments.

Furthermore, although various studies have investigated the effects of NSCs on the regulation of photosynthetic genes in various plants (Sheen, 1994; Van Oosten and Besford, 1995; Moore et al., 1999), only a few comprehensive attempts have been made to elucidate the whole mechanism of NSCs-induced photosynthetic downregulation with transcriptome analysis focusing on both the photosynthetic induction and steady-state photosynthesis (Marquardt et al., 2021). We considered that it is possible to induce NSCs accumulation without reducing N supply in the source leaves by cold girdling, which will clarify either NSCs accumulation or N deficiency could cause the photosynthetic downregulation more directly. Localised cooling was achieved by cold water flowing from a refrigerated circulator to plastic tubing fitted to stems or petioles. It is suggested that viscosity of phloem sap increases and/or sieve plate pores are plugged by localized cooling, which inhibit translocation of photosynthates but not water, inorganic solutes, and amino acids (Peuke et al., 2006). This treatment successfully inhibited phloem transport capacity, resulting in the NSCs accumulation in source leaves of sugar beet (*Beta vulgaris*) (Geiger et al., 1973), tobacco (*Nicotiana tabacum*) (Krapp et al., 1993), common bean (*Phaseolus vulgaris*) (Hannah et al., 2001), and sugarcane (*Saccharum officinarum*) (McCormick et al., 2008a). Sucrose feeding to plants is another approach to induce NSCs accumulation in source leaves (Furbank et al., 1997; Araya et al., 2006). Thus, these methods could be used to accumulate NSCs without affecting N supply in the source leaves.

This study aimed at elucidating either NSCs accumulation or N deficiency contributes to the photosynthetic downregulation more directly in soybean (*Glycine max*) and common bean (*Phaseolus vulgaris*) and how NSCs accumulation affects each of the photosynthetic induction process at the physiological and transcriptomic levels. For the first purpose, we tried to develop a new and simple cold-girdling system with a Peltier device since the previous system with plastic tubing and refrigerated circulator is space consuming, which made it difficult to perform the cold-girdling for many plants at the same time in the environment-controlled growth chamber (Fig. S1). For the second purpose, we evaluated effect of NSCs accumulation triggered by cold-girdling, sucrose feeding, and low N treatments on the photosynthetic induction, steady-state maximum photosynthesis (Fig. S2), and underlying gene expression profiles in *G. max*. We will further discuss regulatory mechanisms of NSCs accumulation and the decrease in leaf N content caused by sink-source imbalance focusing on the nitrogen assimilation/remobilization and sugar transport between chloroplast and cytosol.

## Results

### Response of A and g_s_ to the step change in light intensity

CG, Suc, and LN treatments caused the photosynthetic downregulation by decreasing *A*_max_ and limiting stomatal opening during the photosynthetic induction in *G. max* (Fig. 1), whereas CG treatment did not cause the downregulation in *P. vulgaris* (Fig. 2). In Ct plants of both species and GP plants of *P. vulgaris, A* increased rapidly and *g*_s_ was constantly high during the 20 min high light phase from 0 to 7−8 DAT. In *G. max*, CG plants showed the lowest *A*_max_ and *g*_smax_ and highest *T*_90A_ and *T*_90gs_ from 5 DAT. Meanwhile, *A*_max_ and *g*_smax_ decreased and *T*_90A_ and *T*_90gs_ increased in Suc and LN plants from 7−8 DAT (Fig. S3a, c, Table 1). In *P. vulgaris*, only Suc and LN treatments significantly decreased *A*_max_ and *g*_smax_ and increased *T*_90A_ and *T*_90gs_ from 5−6 DAT (Fig. S3b, d, Table 1). These results were consistent through Exp. 1, 2, and 3 in *G. max* (Fig. S4 a-d).

**Table 1.**
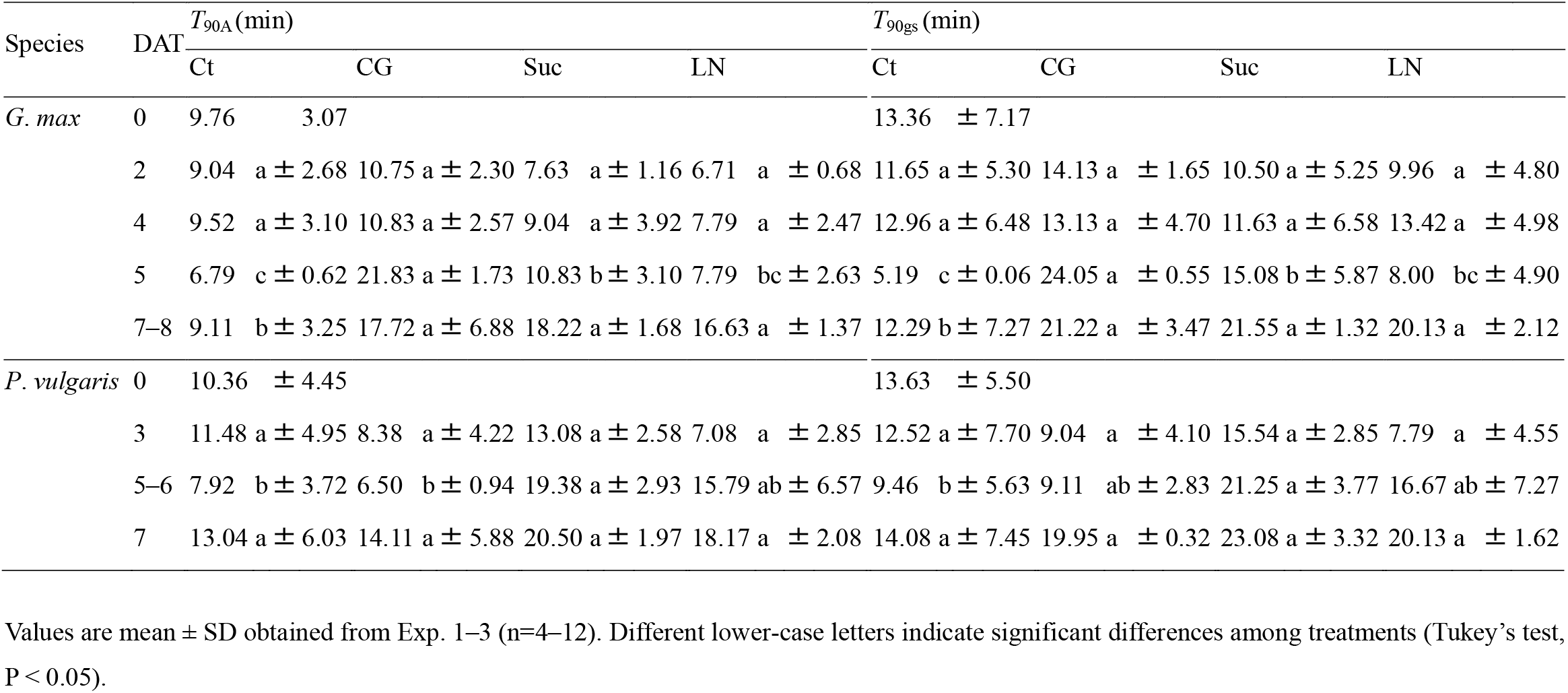
Time at net CO_2_ assimilation (*A*) and stomatal conductance (*g*_s_) reached 90% of the maximum *A* (*A*_max_) and *g*_s_ (*g*_smax_) (*T*_90A_ and *T*_90gs_) in *G. max* and *P. vulgaris*.

**Fig. 1.**
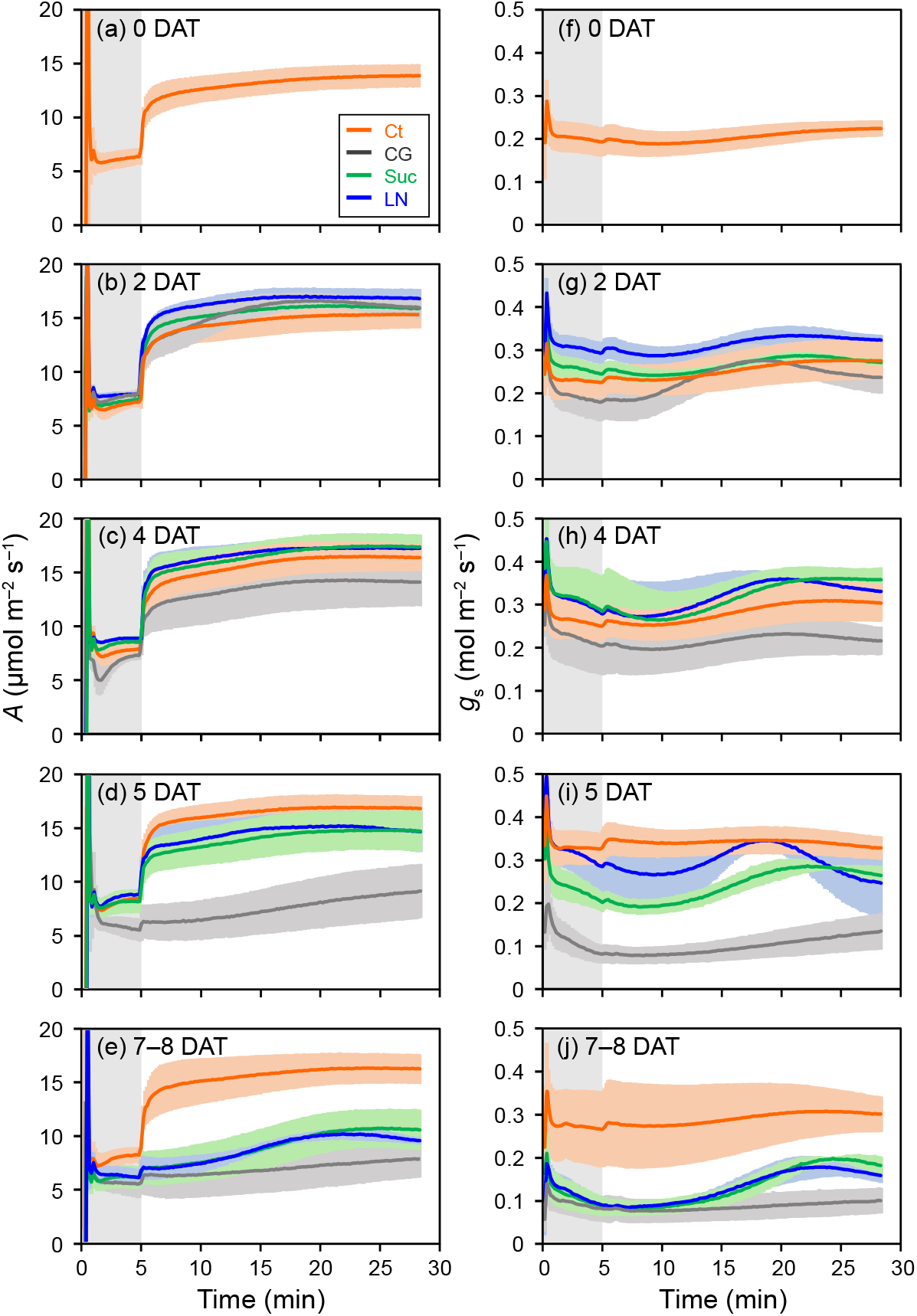
Temporal response of net CO_2_ assimilation (*A*) and stomatal conductance (*g*_s_) to a step change in PPFD from 200 (shaded area) to 1000 (unshaded area) μmol m^−2^ s^−1^ in *G. max*. (a, f) 0 day after treatments (DAT), (b, g) 2 DAT, (c, h) 4 DAT, (d, i) 5 DAT, (e, j) 7–8 DAT. Orange, grey, green, and blue lines indicate control (Ct), cold-girdling (CG), sucrose feeding (Suc), and low nitrogen (LN) treatments, respectively. Values are mean ± SD with a measurement interval of 10 seconds obtained from Exp. 1–3 (n=4–12).

**Fig. 2.**
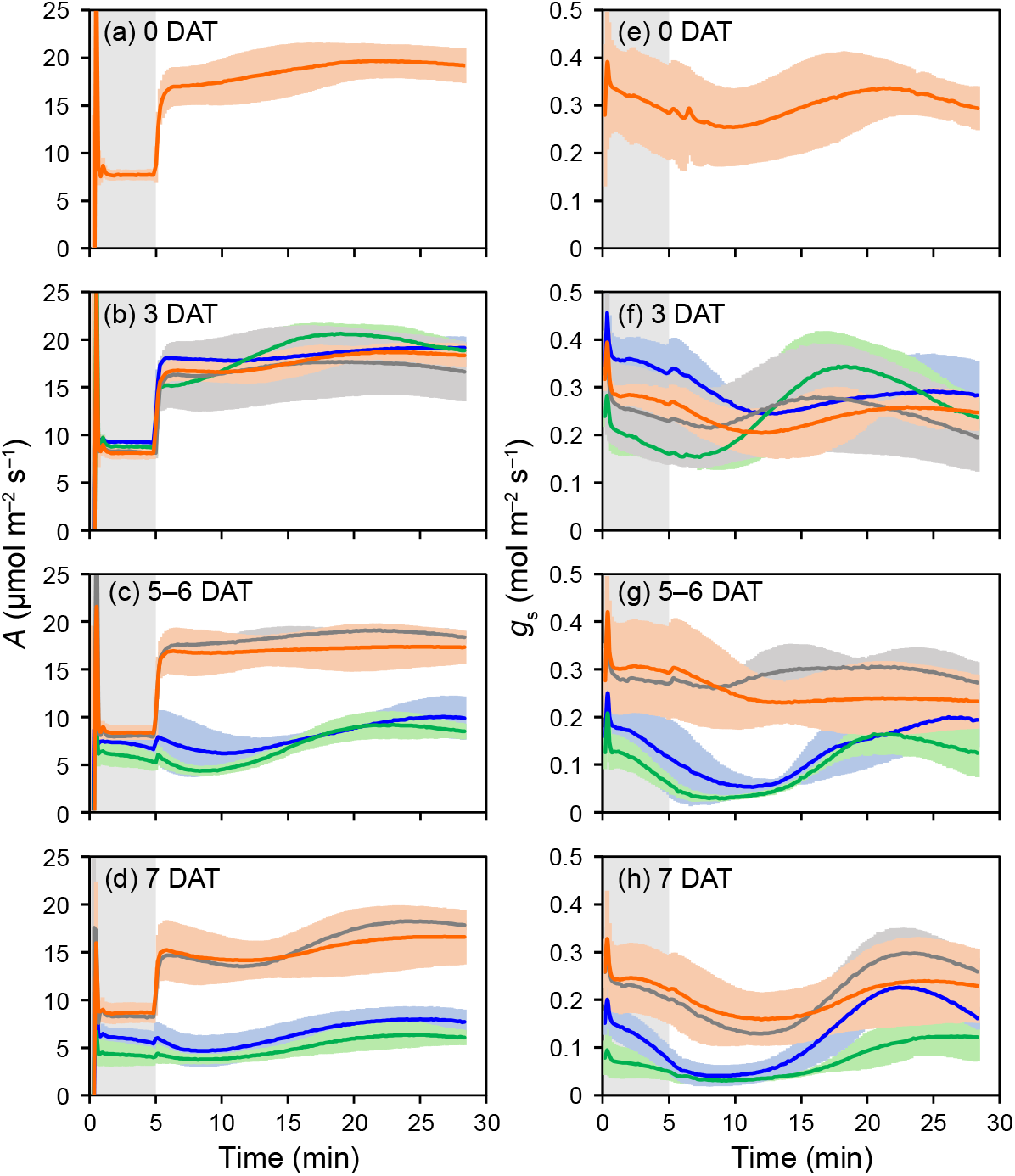
Temporal response of net CO_2_ assimilation (*A*) and stomatal conductance (*g*_s_) to a step change in PPFD from 200 (shaded area) to 1000 (unshaded area) μmol m^−2^ s^−1^ in *P. vulgaris*. (a, e) 0 day after treatments (DAT), (b, f) 2 DAT, (c, g) 5∼6 DAT, (d, h) 7 DAT. Orange, grey, green, and blue lines indicate control (Ct), cold-girdling (CG), sucrose feeding (Suc), and low nitrogen (LN) treatments, respectively. Values are mean ± SD with a measurement interval of 10 seconds obtained from Exp. 1–2 (n=4–8).

### CG, Suc, and LN treatments induced NSCs accumulation and decrease in nitrogen content

In both species, NSCs was mostly composed of starch and the contents of glucose and sucrose were minor, consistent with the previous study (Sugiura et al., 2019). In *G. max*, all experimental treatments aimed at NSCs accumulation (Fig. S1) induced NSCs accumulation in the primary leaves, and CG plants showed the highest NSC_area_ (Fig. 3a). CG, Suc, and LN plants showed significantly lower N_area_ than Ct plants, and LN plants showed the lowest N_area_ (Fig. 3c). Meanwhile, in *P. vulgaris*, CG treatment did not induce NSCs accumulation (Fig. 3b), and N_area_ was significantly lower in Suc and LN plants (Fig. 3d). These results were consistent through Exp. 1, 2, and 3 in *G. max* (Fig. S4 e, f).

**Fig. 3.**
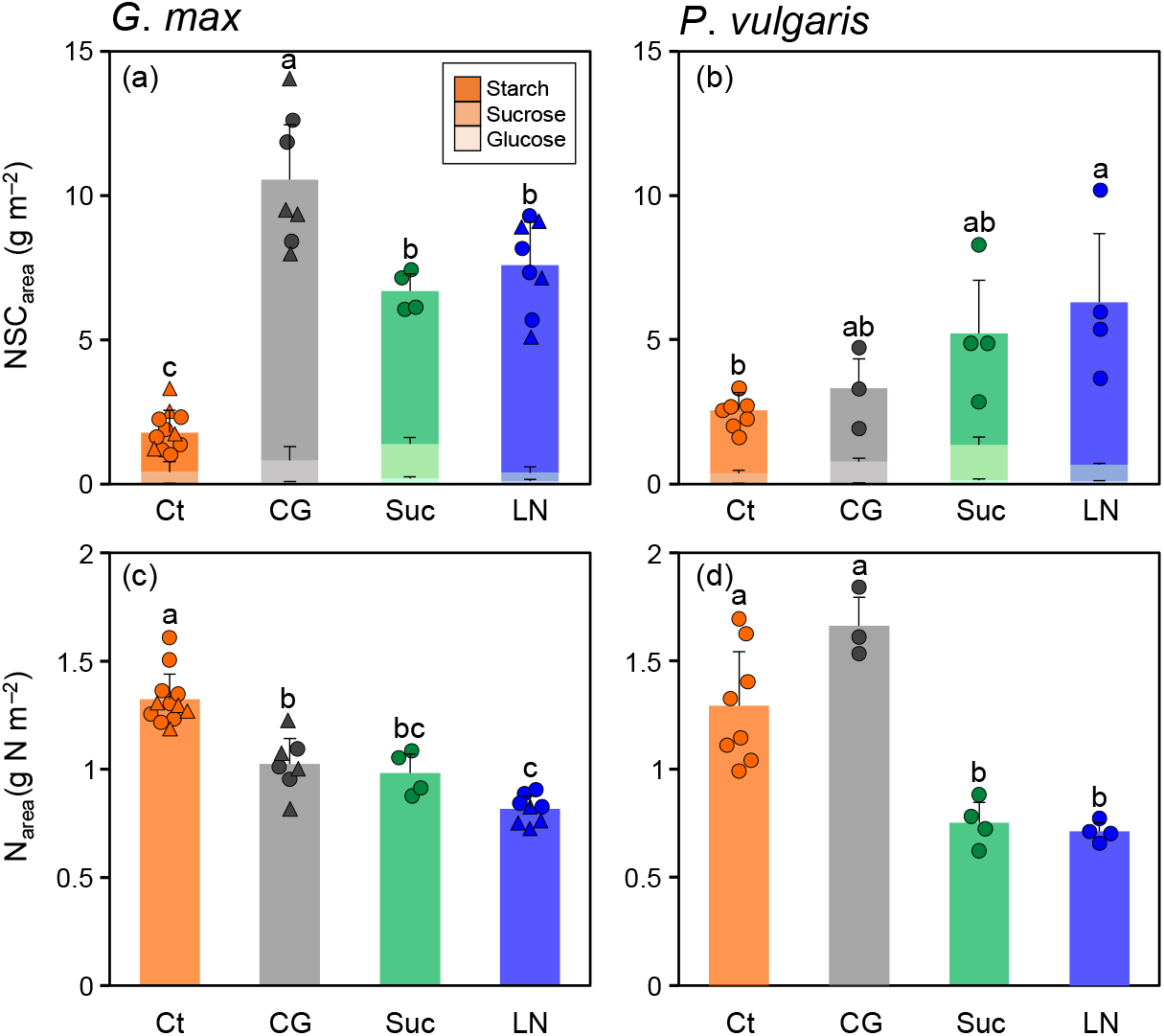
Nonstructural carbohydrate (NSC_area_) and leaf nitrogen content per area (N_area_) in the primary leaves of *G. max* and *P. vulgaris* on 7–8 days after treatments (DAT). (a, c) *G. max* and (b, d) *P. vulgaris*. Orange, grey, green, and blue bar indicate control (Ct), cold-girdling (CG), sucrose feeding (Suc), and low nitrogen (LN) treatments, respectively. NSC_area_ is expressed as sum of starch, sucrose, and glucose. Triangle represents data of *G. max* in Exp. 3. Bars are mean + SD obtained from Exp. 1–3 (n=4–12). Different lower-case letters indicate significant differences among treatments (Tukey’s test, P < 0.05).

In both species of Suc and LN plants, NSCs accumulated not only in the primary leaves but also all other organs (upper leaves, stems, and roots) (Table. 2). Meanwhile, in *G. max*, CG plants showed higher NSCs levels in the upper leaves and stems, but lower NSCs levels in the roots. CG plants of *P. vulgaris* showed similar NSCs levels as Ct plants in all organs.

**Table 2.** Nonstructural carbohydrates (NSCs) in each organ in *G. max* and *P. vulgaris*.

| Species | organs | NSC concentration (mg g<sup>-1</sup>) |  |  |  |  |  |  |  |
| --- | --- | --- | --- | --- | --- | --- | --- | --- | --- |
|  |  | Ct |  | CG |  | Suc |  | LN |  |
| *G. max* | Primary leaves | 62.65 | c ± 21.0 | 279.4 | a ± 35.8 | 198.7 | b ± 22.5 | 236.7 | ab ± 21.1 |
|  | Upper leaves | 37.47 | b ± 17.6 | 135.8 | a ± 10.7 | 128.9 | a ± 9.37 | 154.8 | a ± 7.46 |
|  | Stems | 13.95 | b ± 6.44 | 55.62 | a ± 3.09 | 64.30 | a ± 12.9 | 62.05 | a ± 3.55 |
|  | Roots | 4.953 | b ± 2.65 | 2.582 | b ± 0.267 | 10.92 | a ± 3.74 | 12.72 | a ± 1.51 |
|  | Whole-plant | 30.16 | c ± 9.05 | 130.7 | a ± 15.6 | 103.8 | b ± 10.67 | 117.8 | ab ± 6.39 |
| *P. vulgaris* | Primary leaves | 90.12 | c ± 15.7 | 93.45 | bc ± 22.1 | 172.3 | ab ± 49.7 | 206.6 | a ± 50.3 |
|  | Upper leaves | 79.87 | b ± 28.6 | 81.66 | b ± 48.0 | 190.1 | a ± 16.3 | 223.8 | a ± 19.9 |
|  | Stems | 38.62 | b ± 13.7 | 39.05 | b ± 2.46 | 142.5 | a ± 6.92 | 132.6 | a ± 11.8 |
|  | Roots | 8.959 | b ± 3.88 | 5.518 | b ± 1.52 | 25.66 | a ± 4.62 | 19.04 | a ± 3.97 |
|  | Whole-plant | 60.32 | b ± 12.7 | 66.94 | b ± 8.79 | 129.6 | a ± 11.6 | 140.3 | a ± 24.0 |

NSCs was determined from plants harvested on the last day of each experiment. Values are mean ± SD obtained from Exp. 1–3 (n=4–12). Different lower-case letters indicate significant differences among treatments (Tukey's test, P < 0.05).

### CG, Suc, and LN treatments did not affect whole-plant growth but biomass allocation

In *G. max*, N_mass_ was decreased by all experimental treatments, but Suc and LN plants showed significantly lower N_mass_ than CG plants at the whole-plant level (Table 3). In *P. vulgaris*, N_mass_ was decreased by Suc and LN treatments, but CG treatment did not induce the decrease in N_mass_.

**Table 3.** Nitrogen content (N_mass_, g N g^−1^) in the leaves, stems, and roots in *G. max* and *P. vulgaris*.

| Species | organs | $N_{\text{mass}}$ (mg N g<sup>-1</sup>) | | | | | | | | | |
| --- | --- | --- | --- | --- | --- | --- | --- | --- | --- | --- | --- |
|  |  | Ct |  | CG |  | Suc |  | LN |  |  |  |
| *G. max* | Primary leaves | 53.63 | a ± 3.21 | 26.25 | b ± 2.95 | 29.05 | b ± 2.58 | 27.20 | b ± 1.52 |  |  |
|  | Upper leaves | 60.86 | a ± 1.56 | 40.34 | b ± 2.16 | 34.48 | c ± 1.11 | 31.26 | c ± 1.86 |  |  |
|  | Stems | 35.81 | a ± 2.05 | 22.54 | b ± 1.01 | 20.80 | b ± 2.57 | 20.42 | b ± 0.94 |  |  |
|  | Roots | 47.31 | a ± 3.46 | 50.69 | a ± 1.62 | 28.29 | b ± 0.62 | 21.51 | c ± 0.72 |  |  |
|  | Whole-plant | 49.87 | a ± 1.82 | 33.31 | b ± 1.93 | 28.16 | c ± 1.05 | 25.18 | c ± 0.92 |  |  |
| *P. vulgaris* | Primary leaves | 45.33 | a ± 3.96 | 48.35 | a ± 2.13 | 25.39 | b ± 1.29 | 24.26 | b ± 1.80 |  |  |
|  | Upper leaves | 57.00 | a ± 5.05 | 57.43 | a ± 6.60 | 31.63 | b ± 2.21 | 25.36 | b ± 1.49 |  |  |
|  | Stems | 27.94 | b ± 3.75 | 35.08 | a ± 3.91 | 13.04 | c ± 1.36 | 10.93 | c ± 0.64 |  |  |
|  | Roots | 39.75 | b ± 3.23 | 45.13 | a ± 1.82 | 26.90 | c ± 1.78 | 23.30 | c ± 1.06 |  |  |
|  | Whole-plant | 44.43 | a ± 3.01 | 46.28 | a ± 1.59 | 24.66 | b ± 1.45 | 21.43 | b ± 0.97 |  |  |

$N_{\text{mass}}$ was determined from plants harvested on the last day of each experiment. Values are mean ± SD obtained from Exp. 1–3 (n=4–12). Different lower-case letters indicate significant differences among treatments (Tukey's test, $P < 0.05$ ).

In *G. max*, CG plants showed the highest LMA, and Suc and LN plants also showed higher LMA than Ct plants, whereas in *P. vulgaris*, LMA was not different among all plants (Table 4). In both species, total biomass was not affected by all experimental treatments. In both species, Suc and LN treatments decreased leaf-to-root ratio compared with Ct plants, while CG treatment increased leaf-to-root ratio in *G. max*.

**Table 4.** Leaf mass per area (LMA) of the primary leaves, total biomass, and leaf-to-root ratio in *G. max* and *P. vulgaris*.

|  | Species | Treatment |  |  |  |  |  |  |  |
| --- | --- | --- | --- | --- | --- | --- | --- | --- | --- |
|  |  | Ct |  | CG |  | Suc |  | LN |  |
| LMA | *G. max* | 25.45 | c ± 2.06 | 42.60 | a ± 2.12 | 33.80 | b ± 1.66 | 31.16 | b ± 3.08 |
| (g m<sup>-2</sup>) | *P. vulgaris* | 24.41 | a ± 1.93 | 25.91 | a ± 3.14 | 29.62 | a ± 3.36 | 29.58 | a ± 3.99 |
| Total biomass | *G. max* | 0.697 | a ± 0.22 | 0.716 | a ± 0.047 | 0.769 | a ± 0.21 | 0.706 | a ± 0.12 |
| (g) | *P. vulgaris* | 1.17 | a ± 0.23 | 1.02 | a ± 0.033 | 0.987 | a ± 0.21 | 1.06 | a ± 0.20 |
| Leaf to root ratio | *G. max* | 3.09 | b ± 0.32 | 4.99 | a ± 0.43 | 2.50 | bc ± 0.29 | 2.21 | c ± 0.12 |
| (g g<sup>-1</sup>) | *P. vulgaris* | 2.80 | a ± 0.23 | 2.82 | a ± 0.085 | 1.78 | b ± 0.20 | 1.72 | b ± 0.24 |

These traits were determined from plants harvested on the last day of each experiment. Values are mean ± SD obtained from Exp. 1–3 (n=4–12). Different lower-case letters indicate significant differences among treatments (Tukey's test, P < 0.05).

### Relationships between photosynthetic traits, NSC, and leaf nitrogen

*A*_max_ and *g*_smax_ were strongly negatively correlated with NSC_area_ in *G. max*, (Fig. 4a, b), whereas weaker negative correlations were observed in *P. vulgaris* (Fig. S5a, b). *A*_max_ was strongly positively correlated with *g*_smax_ and N_area_ in both species (Figs. 4c, d, S5c, d). In *G. max*, CG plants showed lower *A*_max_ relative to N_area_ (Fig. 4d).

**Fig. 4.**
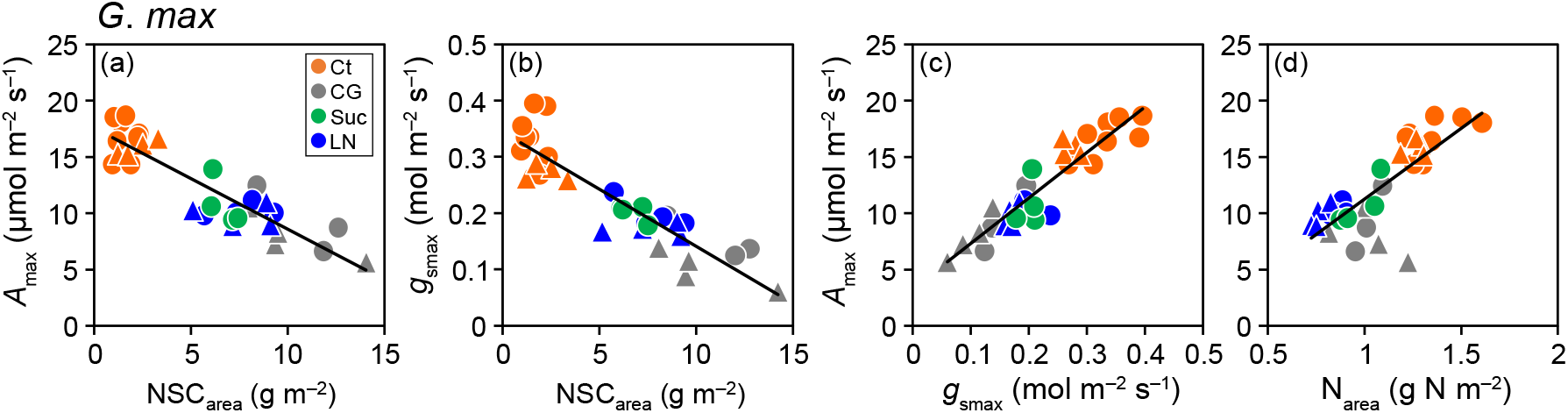
Relationships between photosynthetic traits (*A*_max_ and *g*_smax_), nonstructural carbohydrate (NSC_area_), and leaf nitrogen content per area (N_area_) in the primary leaves of *G. max* on 7–8 days after treatments (DAT). Orange, grey, green, and blue symbols indicate control (Ct), cold-girdling (CG), sucrose feeding (Suc), and low nitrogen (LN) treatments, respectively. Triangle represents data of *G. max* in Exp. 3. Values of R^2^ are (a) 0.77, (b) 0.71, (c) 0.86, (d) 0.61 (*P* < 0.01).

### Transcriptome of primary leaves in response to CG and LN treatments

In Exp. 3, RNA-seq analysis of the primary leaves revealed changes in gene expression profiles in response to CG and LN treatments in *G. max*. Principal component analysis (PCA) classified the RNA-Seq data into three groups according to the treatments (Fig. S6). The K-means clustering analysis revealed the three gene clusters (A, B, C) with either increased or decreased expression in response to the treatments and one cluster (D) with unclear characteristics (Fig. 5, Table S1). Characterization of each cluster by pathway enrichment analysis showed that cluster B which is downregulated in CG and LN plants was enriched in Gene Ontology (GO) terms associated with photosynthesis. Cluster A which is upregulated in CG included GO terms associated with oxidative and temperature stresses, whereas Cluster C which is upregulated in LN included GO terms associated with cell wall metabolisms. Information about normalized transcript levels of genes in each cluster was provided in Table S2.

**Fig. 5.**
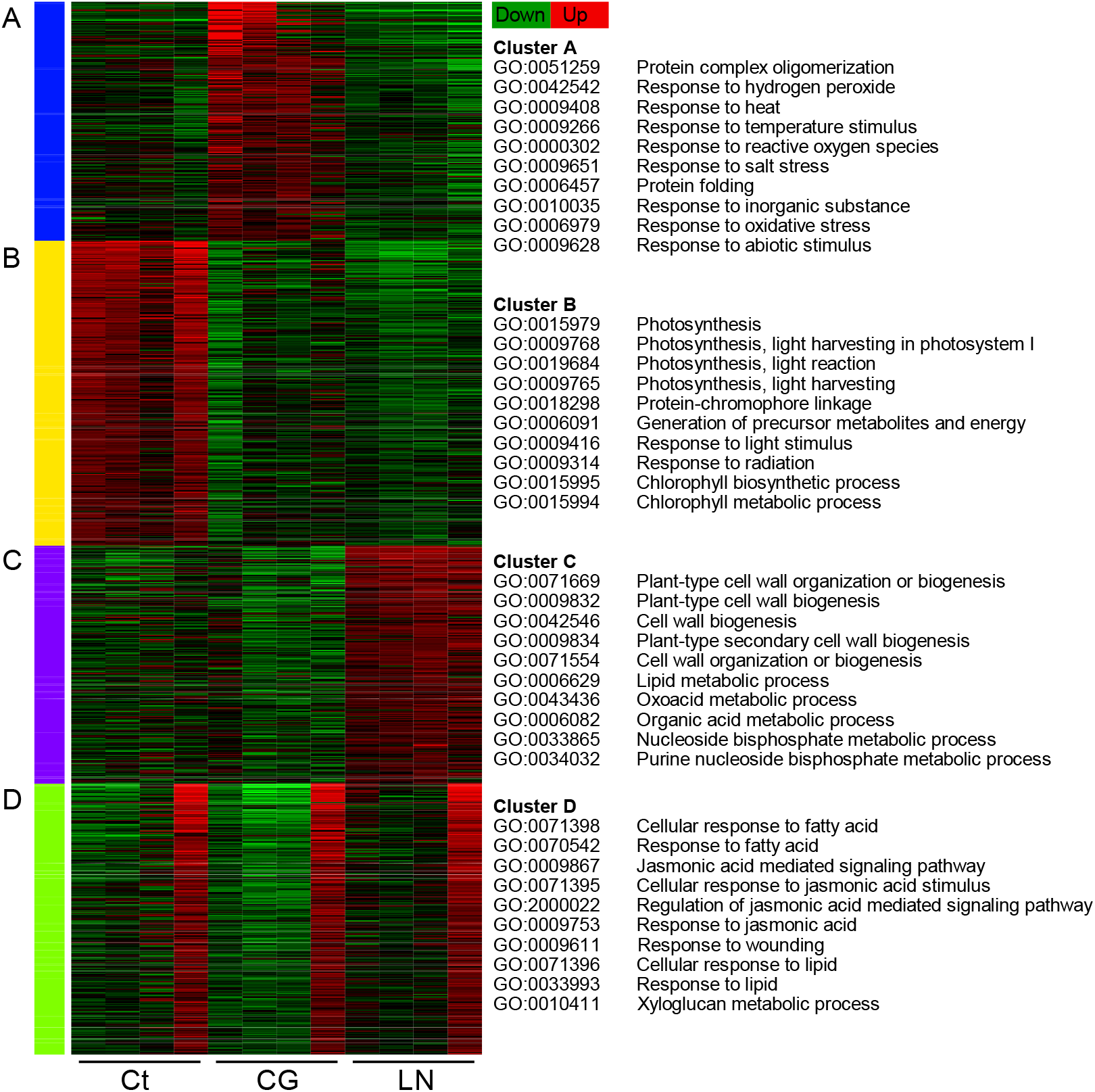
K-means clustering and gene ontology (GO) enrichment analysis from transcriptome data of *G. max* obtained in Exp. 3. Heat map shows up-(red) and down-(green) regulated genes in control (Ct), cold-girdling (CG), and low nitrogen (LN) (n = 4). 2000 genes were grouped into 4 clusters, each of which include 442 (Cluster A), 557 (Cluster B), 473 (Cluster C), and 508 genes (Cluster D) (Supplementary Table X). Top 10 GO biological processes were shown for each cluster.

### Calvin cycle genes were downregulated in response to sugar accumulation

Analysis of photosynthetic genes showed that the transcript levels of *RbcS* encoding a small subunit of Rubisco as well as other key enzymes including PRK (Phosphoribulokinase) that regenerate RuBP were downregulated in CG and LN plants (Fig. 6, Table S3) with lower *A*_smax_ (Figs. 1, S3). Furthermore, the expression of Rubisco activase (*Rca*) that facilitates Rubisco activation (Portis Jr et al., 2008) and Rubisco accumulation factor (*RAF*) that is required for assembly and stability of Rubisco (Fristedt et al., 2018) was also downregulated. Expression genes involved in photochemical reaction, such as subunits and light harvesting complexes of photosystem I and II, *Psa, Psb*, and *LHC* (Tables S4, 5) as well as cytochrome b_6_f complexes such as *Pet* and *FD* (Table S6), were downregulated in CG and LN plants.

**Fig. 6.**
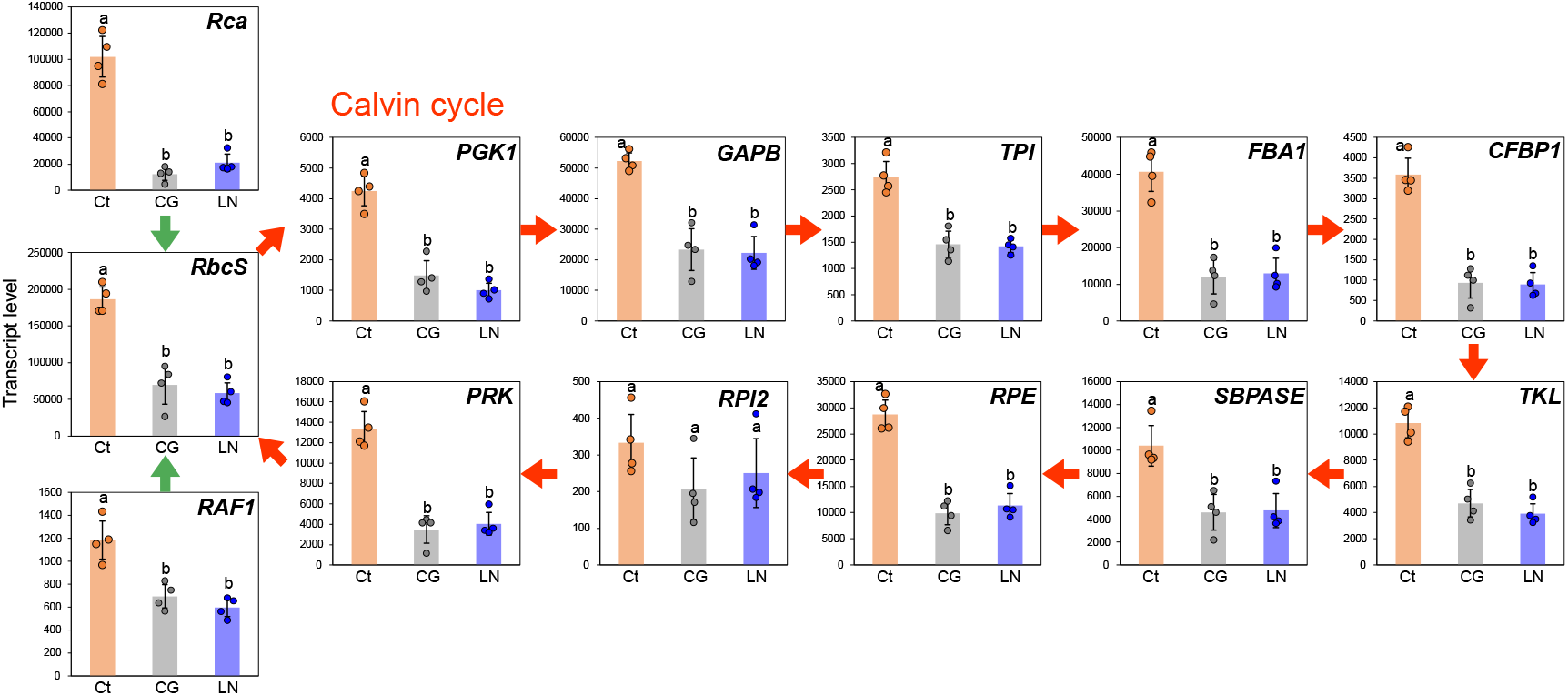
Expression of key genes in the Calvin cycle in response to cold-girdling (CG) and low nitrogen (LN). Data are obtained from transcriptome data of *G. max* in Exp. 3. Transcript levels of genes annotated with Arabidopsis gene ID are calculated as the total read counts of duplicated genes of *G. max* (Supplementary Table 3). RCA is Rubisco activase that facilitates Rubisco activation, and RAF is Rubisco accumulation factor required for assembly and stability of Rubisco. PRK is responsible for the regeneration of RuBP. Bars are mean± SD (n=4). Different lower-case letters indicate significant differences among treatments (Tukey’s test, P < 0.05).

### Genes promoting stomatal opening were downregulated in response to sugar accumulation

The gene expression profiles in guard cells showed that the transcript levels of factors promoting stomatal opening rather than stomatal closure is downregulated in CG and LN plants (Fig. 7, Table S7) with lower *g*_smax_ and slower stomatal opening during the photosynthetic induction (Figs. 1, S3). We found that *PHOT1* responsible for the blue light dependent stomatal opening (Kinoshita et al., 2001) was significantly downregulated in CG and LN plants. Among the identified three H^+^-ATPase (*AHA2, 5, 11*) facilitating stomatal opening through the hyperpolarization of the plasma membrane (Kimura et al., 2015), only *AHA11* was significantly downregulated in CG compared with Ct plants, and *PATROL1* involved in the localization of H^+^-ATPase (Hashimoto-Sugimoto et al., 2013) was unaffected by CG and LN treatments. We also found the transcript level of *KAT1*, an inward potassium channel preferentially expressed in guard cells (Nakamura et al., 1995), was significantly decreased by CG and LN treatments.

**Fig. 7.**
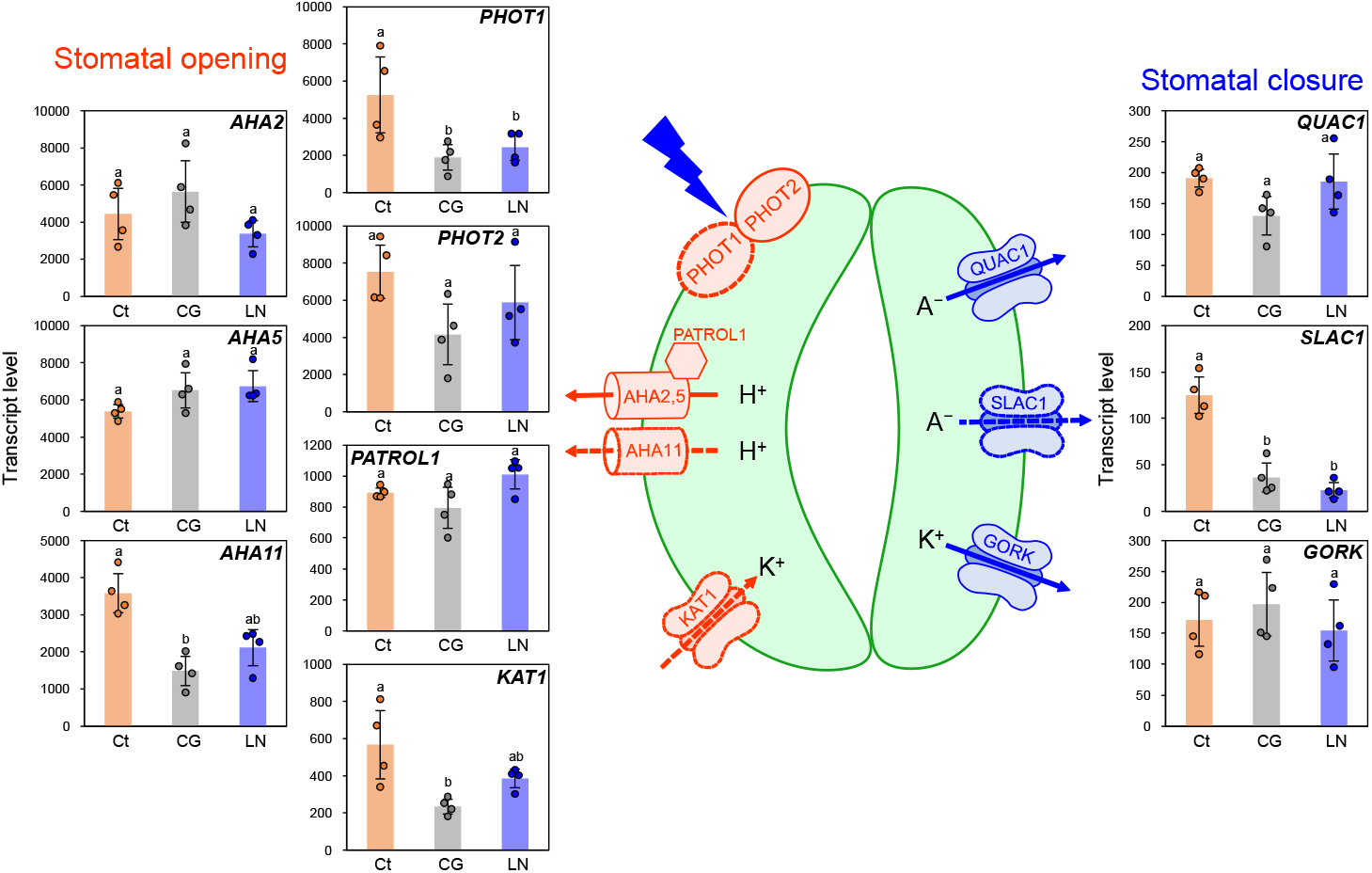
Changes in expression of genes promoting stomatal opening or closure in response to cold-girdling (CG) and low nitrogen (LN). Data are obtained from transcriptome data of *G. max* in Exp. 3. Transcript levels of genes annotated with Arabidopsis gene ID are calculated as the total read counts of duplicated genes of *G. max* (Supplementary Table 7). PHOT1 and PHOT2 are blue-light receptors, and PATROL1 is involved in the localization of H^+^-ATPase. KAT1 and GORK are inward and outward potassium (K^+^) channels, respectively, and QUAC1 and SLAC1 are outward anion (A^−^) channels. Bars are mean ± SD (n=4). Different lower-case letters indicate significant differences among treatments (Tukey’s test, P < 0.05).

Although we explored genes promoting stomatal closure such as *GORK* (Hosy et al., 2003) and *QUAC* (Imes et al., 2013), no candidates were found to be upregulated and only *SLAC1* which is important for normal stomatal closure (Vahisalu et al., 2008) was downregulated in CG and LN plants.

### Nitrogen assimilation and remobilization in response to sugar accumulation

Since leaf nitrogen content was decreased even by CG treatment, we hypothesized that nitrogen metabolism genes would be affected by the accumulation of NSCs. As hypothesized, transcript levels of genes involved in the nitrogen transport and assimilation were downregulated while those involved in the nitrogen remobilization were upregulated by the accumulation of NSCs (Fig. 8, Table S8). Among nitrate transporters identified, NRT1.1, a dual affinity nitrate transporter expressed also in shoots (Ho et al., 2009), was significantly downregulated in CG and LN. Except for cytosolic glutamine synthase (GLN1.1), all the gene expressions involved in the nitrogen assimilation such as nitrate and nitrite reductases (NR and NiR), plastidic glutamine synthase (GLN2), and glutamate synthase (FD-GOGAT) were significantly decreased in CG and LN plants. Meanwhile, transcript levels of genes responsible for source-to-sink nitrogen remobilization such as NRT1.7 facilitating phloem loading of nitrate (Fan et al., 2009) and cationic and neutral amino acids transporters (CAT and AAP) (Liu and Bush, 2006; Zhou et al., 2020; Liang et al., 2022) were significantly upregulated in CG and LN plants.

**Fig. 8.**
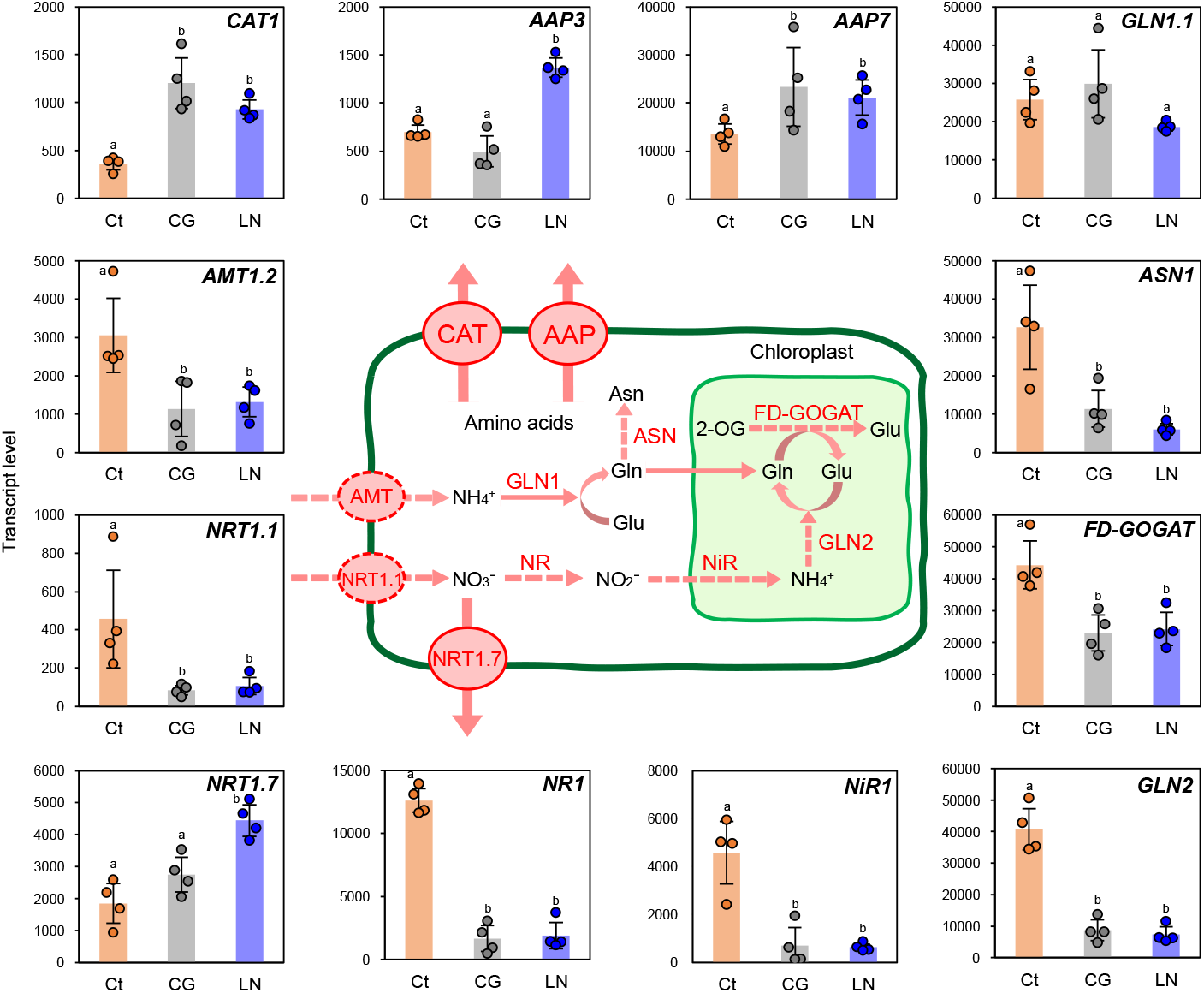
Changes in expression of gene involved in nitrogen assimilation in response to cold-girdling (CG) and low nitrogen (LN). Data are obtained from transcriptome data of *G. max* in Exp. 3. Transcript levels of genes annotated with Arabidopsis gene ID are calculated as the total read counts of duplicated genes of *G. max* (Supplementary Table 8). NRT and AMT are NO_3_^−^ and NH_4_^+^ transporter, respectively, where NRT1.7 is responsible for phloem loading of NO_3_^−^. NR and NiR are nitrate and nitrite reductase, respectively. GLN, FD-GOGAT, and ASN are glutamine (Gln), glutamate (Glu), and asparagine (Asn) synthase, respectively. CAT and AAP are cationic and neutral amino acid transporters, respectively. Bars are mean± SD (n=4). Different lower-case letters indicate significant differences among treatments (Tukey’s test, P < 0.05).

### Sugar transport/metabolism rather than starch synthesis/degradation influenced NSCs accumulation

We further investigated the gene expression profiles underlying NSCs accumulation in the source leaves focusing on the starch and sugar metabolism in the chloroplast and cytosol (Bahaji et al., 2014). The results showed that starch synthesis and degradation pathways in the chloroplast were not affected, while hexose-phosphate translocators in the chloroplast membrane were directed to transport hexose-phosphate into the chloroplasts in CG and LN plants (Fig. 9). Although we expected gene expression of enzymes involved in starch synthesis would be upregulated in CG and LN plants, key genes such as *APL* and *APS* encoding the subunits of AGPase and *SS* encoding starch synthase remained mostly unchanged (Table S9) as well as genes in starch degradation pathway (Table S10). On the other hand, transcript levels of genes encoding the triose phosphate (TP)/phosphate (Pi) translocator (*TPT*) that exports TP in exchange for inorganic phosphate (Weise et al., 2019) were significantly downregulated, while transcript levels of genes encoding the glucose-6-phosphate (G6P)/phosphate translocator 1 and 2 (*GPT1, GPT2*) that imports G6P to the chloroplast (Dyson et al., 2015) were significantly upregulated especially in CG plants (Table S11). Among sucrose-phosphatase (SPP) and sucrose-phosphate synthase (SS) which play a major role in sucrose synthesis (Lunn and MacRae, 2003), *SPS3* was downregulated in CG and LN plants, and gene expression of *SUT1* and *SUT2* that play a major role in sucrose transport from source to sink organs were not altered (Table S9). Transcript levels of Hexokainase (*HXK1, HXK2, HXK3*), which phosphorylate hexoses and are proposed to be functioning as a sugar sensor (Jang et al., 1997; Moore et al., 1999), and *SnRK1*, which is also indicated as another sugar sensor (Baena-González et al., 2007), did not differ among all plants except that expression level of *HXK3* was higher in LN plants (Table S12).

**Fig. 9.**
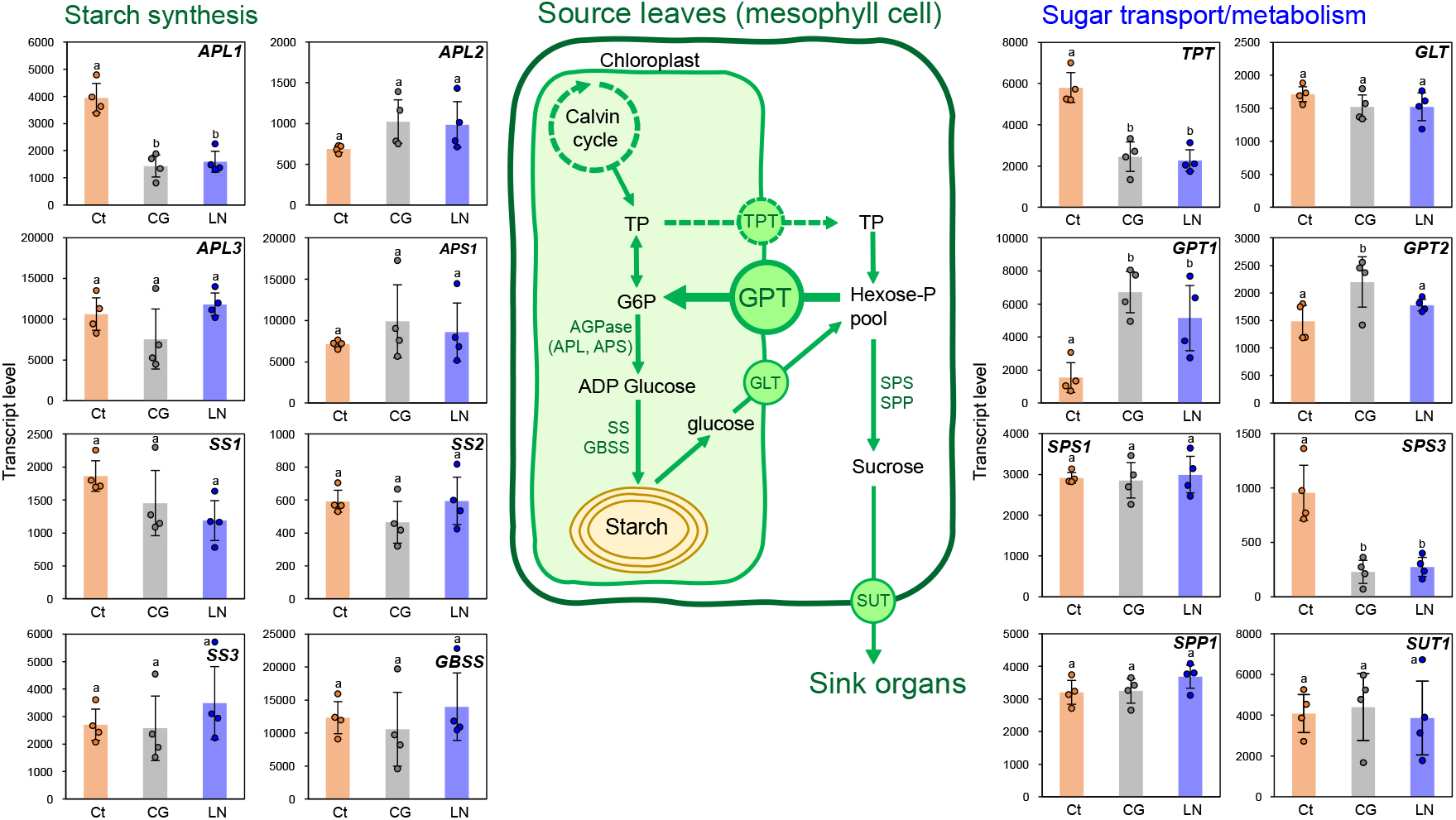
Changes in expression of gene involved in starch synthesis and sugar transport/metabolism in response to cold-girdling (CG) and low nitrogen (LN). Data are obtained from transcriptome data of *G. max* in Exp. 3. Transcript levels of genes annotated with Arabidopsis gene ID are calculated as the total read counts of duplicated genes of *G. max* (Supplementary Tables 9-11). APL and APS are large and small subunits of ADP glucose pyrophosphorylase (AGPase), respectively. SS and GBSS are soluble and granule bound starch synthases, respectively. TPT and GLT export triose phosphate (TP) and glucose from the chloroplast to the cytosol, respectively. GPT transport glucose-6-phosphate (G6P) from the cytosol to the chloroplast. SPS and SPP play a major role in sucrose synthesis, and SUT is sucrose transporter. Bars are mean± SD (n=4). Different lower-case letters indicate significant differences among treatments (Tukey’s test, P < 0.05).

## Discussion

### NSCs accumulation directly triggered the photosynthetic downregulation

In the present study, we established an experimental system for inducing NSCs accumulation without reducing N supply in the source leaves, which revealed that NSCs accumulation rather than N deficiency directly decreased maximum photosynthetic capacity and delayed photosynthetic induction in *G. max*. Our comprehensive transcriptomic analysis further revealed the coordinated downregulation of genes involved in the three steps of photosynthetic induction process: (1) activation of the RuBP regeneration process, (2) activation of Rubisco, and (3) stomatal opening. These results provided an overall picture of the photosynthetic downregulation and NSCs accumulation driven by changes in sink-source balance.

### Cold-girdling with the Peltier device is an effective method for inducing NSCs accumulation

Our cold-girdling technique using the Peltier device, which requires less space and efforts compared to the previous system using a refrigerated circulator, successfully induced the highest NSCs accumulation in CG plants of *G. max* (Fig. 3a). It is remarkable that NSCs level was higher in the stems and upper leaves while it was lower in the roots (Table2), demonstrating that phloem transport was clearly inhibited, and sink-limited situations was created by cooling stems with the Peltier device in *G. max*. The results also showed that exogeneous sucrose feeding can induce similar level of NSCs accumulation to LN treatment in *G. max* as reported in *P. vulgaris* (Araya et al., 2006; Sugiura et al., 2019), confirming the validity of these method for inducing NSCs accumulation.

It also should be noted that CG treatment could not suppress the phloem sugar transport in *P. vulgaris*. Hannah et al. (2001) reported a large genotypic difference in the effect of cold-girdling on suppressing phloem transport in *P. vulgaris*, which may be due to differences in structural and physiological characteristics of the sieve tubes, such as callose and P-proteins plugging sieve tubes (Knoblauch et al., 2014). Nevertheless, once validated in a specific plant species, the present cold-girdling system can work as a powerful tool for assessing physiological responses to sugar accumulation and for exploring shoot-to-root signalling with minimum effects on the nitrogen supply and whole-plant growth (Tables 3, 4).

### NSCs accumulation rather than N deficiency directly decreased maximum photosynthetic capacity and delayed photosynthetic induction

Temporal response of *A* and *g*_s_ to the increased PPFD showed that all the treated plants not only decreased *A*_max_ and *g*_smax_ but also increased *T*_90A_ and *T*_90gs_ except for CG plants of *P. vulgaris*, suggesting that NSCs accumulation caused the delayed photosynthetic induction (Figs. 1, 2, Table 1). In *G. max*, since these photosynthetic characteristics were downregulated earlier in CG plants than in LN and Suc plants (Fig. 1d, i), NSCs may have begun to accumulate at 4DAT and reached the highest NSC_area_ at 7−8 DAT (Fig. 3a). This result is consistent with previous studies showing time required for the downregulation of maximum photosynthesis after CG treatment ranges from less than 24 hours in *Saccharum officinarum* (McCormick et al., 2008a), 4 days in *Spinacia oleracea* (Krapp and Stitt, 1995), to 1 week in *P. vulgaris* (Hannah et al., 2001), while this is the first study to report that NSCs accumulation caused the delayed photosynthetic induction due to the downregulation of photosynthetic genes.

The strong correlations among *A*_max_, *g*_smax_, NSC_area_, and N_area_ suggest that NSCs accumulation caused the comprehensive decrease in stomatal conductance and the content of photosynthetic proteins in *G. max* (Fig. 4). However, CG plants showed lower *A*_max_ despite higher *N*_area_ than LN plants (Figs. 3c, 4d, S4), suggesting that NSCs also caused the decreased Rubisco activation state as evidenced by the lower transcript level of *Rca* (Fig. 6), which further caused delayed photosynthetic induction (Tanaka et al., 2019). Lower N_area_ in CG plants compared to Ct plants can be attributed the coordinated downregulation of nitrogen assimilation and upregulation of nitrogen remobilization (Fig. 8), in addition to higher leaf-to-root ratio (lower root mass fraction) due to inhibition of carbon translocation to the roots (Table 4).

However, the decrease in *A*_max_ and *g*_smax_ of CG plants observed at an earlier stage (Fig. 1d, i) suggests that NSCs accumulation would be the primary cause of the photosynthetic downregulation. Furthermore, since there were no marked changes in the transcript levels of senescence-associated genes (*SAGs*) (Zhang et al., 2021) at 7 DAT (Table S13), the altered nitrogen metabolism can be considered as active responses for maximizing nitrogen use efficiency independent of leaf senescence (Masclaux-Daubresse et al., 2008; White et al., 2016; Havé et al., 2017).

The difference between LMA and NSCs defined as structural LMA (Bertin et al., 1999), which strongly correlates with cell wall mass per leaf area (Sugiura et al., 2019), was remarkably increased in CG plants of *G. max* (Table 4). This indicated that a decrease in mesophyll conductance by cell wall thickening also contributed to the photosynthetic downregulation in CG plants of *G*.*max* as reported by Sugiura et al. (2020). Overall, the present experimental design allows use to evaluate NSCs accumulation and N deficiency separately, and the results obtained indicate that the photosynthetic downregulation cannot be explained simply by N deficiency (Paul and Driscoll, 1997; Geiger et al., 1999), but is triggered by NSCs accumulation due to sink-source imbalance (Kasai, 2008; White et al., 2016; Sugiura et al., 2017).

It is reported that the increase in biomass and yield under FACE experiments is far less than expected from the chamber experiments under stable conditions (Leakey et al., 2009). This may be due to the accumulation of NSCs, which delayed photosynthesis induction in addition to decreasing maximum photosynthetic capacity, resulting in decreased biomass production under elevated CO_2_ and light-fluctuating conditions. Through understanding dynamic photosynthesis in a fluctuating light regime at elevated CO_2_ (Kaiser et al., 2017), these issues need to be addressed for improved crop production in the future when NSCs accumulation will be more pronounced due to elevated CO_2_ and low fertilizer application.

### Coordinated downregulation of photosynthetic genes triggered by NSCs

The transcriptome analysis of the photosynthetic pathways revealed reduced transcript levels of genes involved in the Calvin cycle (Fig. 6), photochemical reactions (Tables S4-6), and stomatal opening, indicating that NSCs accumulation induced the downregulation of photosynthetic genes in a coordinated manner. This idea was strongly supported by the clustering and GO enrichment analysis, which showed clear differences in downregulated genes among Ct, CG, and LN plants, with the top 10 pathways being dominated by photosynthesis-related pathways in cluster B (Figure 5). Previous study showed clear consistency between the gene expression levels obtained from transcriptome analysis and quantitative PCR in *A. thaliana* (Monden et al., 2022), which also suggests the robustness of the present gene expression analysis.

While there have been numerous studies on the sugar repression of key Calvin cycle genes (Krapp et al., 1993; Nie et al., 1995; Stitt and Krapp, 1999), few studies have examined how sugar accumulation alters transcriptomic profiles of other key photosynthetic enzymes such as *PRK, Rca*, and *RAF*, and photosystem proteins such as *Psa, Psb*, and *LHC* (Van Oosten and Besford, 1995; Marquardt et al., 2021). Although recent studies have shown a lack of correlation between photosynthetic induction and Rca levels in *Oryza sativa* and *G. max* grown under non-stressed conditions (Soleh et al., 2017; Acevedo-Siaca et al., 2020), the present result suggests that Rca levels are controlled in response to sink-source imbalance. Although we also found several genes that are preferentially expressed in guard cells and appear to play an important role in stomatal opening were downregulated (Fig. 7), it is inconclusive whether the observed gene expression profiles are responsible for the suppression of stomatal opening caused by the accumulation of NSCs in guard cells. This is because starch is a necessary substrate for normal stomatal function, providing the carbon skeleton for malate and producing sucrose, which is essential for K^+^ uptake, osmolytes, or as a respiratory energy source.

(Azoulay-Shemer et al., 2016; Lawson and Matthews, 2020). Another possibility is that mesophyll-derived signals suppressed stomatal opening (Fujita et al., 2013; Fujita et al., 2019). Fujita et al. (2019) clearly showed that mesophyll cells pretreated at elevated CO_2_ suppressed stomatal opening of transplanted epidermal tissue, confirming the presence of substances promoting stomatal closure released from mesophyll cells and transferred to guard cells via apoplasts. Future work to elucidate the mechanisms of stomatal regulation will require a combination of new techniques such as single-cell RNA-seq and metabolomic analysis targeting specific mesophyll tissues and guard cells.

### Balancing the sugar-phosphate translocators promotes starch accumulation in chloroplast in response to sink-source imbalance

Transcriptomic data of starch and sugar metabolism in the chloroplast and cytosol suggests that upregulation of *GPT1* and *GPT2* and downregulation of *TPT* promote the translocation of sugar-phosphate from the cytosol to the chloroplast, leading to the promotion of starch accumulation in the chloroplast of *G. max* (Fig. 9). This idea is supported by the unchanged gene expression profiles in the starch biosynthesis/degradation pathways (Tables S9, S10) and partially downregulated sucrose synthesis pathway (Table S9). In addition to the marked upregulation of *GPT2* expression observed in the cold-girdled leaves of *S. officinarum* (McCormick et al., 2008b) and Arabidopsis mutants with excess sugar accumulation (Lloyd and Zakhleniuk, 2004), previous studies have shown the importance of *GPT* in starch accumulation by transporting G6P to the chloroplasts (Dyson et al., 2015; Weise et al., 2019). The present results further suggest that downregulation of *TPT* also contributes to starch accumulation and hexose phosphorylation by retaining TP in the chloroplast and inorganic phosphate in the cytoplasm in response to sugar signal reflecting sink-source imbalance. Since it is essential for excess sugars to be stored in chloroplasts as starch granules to maintain photosynthetic capacity (Mizokami et al., 2019), tuning the balance of these translocators may allow for the creation of plants that exhibit high photosynthetic rates even under sink-limited or stress conditions.

We could not find marked differences in the expression profiles of *HXKs* and *SnRK1* among Ct, CG, and LN plants of *G. max* (Table S12), both of which has been proposed to be involved in sensing and signalling of sugar in relation to trehalose-6-phosphate (Thompson et al., 2017). Comprehensive analysis of *HXK, SnRK1*, and trehalose-related metabolic pathways with the present experimental design enabling manipulation of sink-source balance will reveal whole picture of the sugar sensing/signalling mechanism and photosynthetic downregulation.

## Conclusion

In the present study, we demonstrated that NSCs accumulation directly decreased the maximum photosynthetic capacity and delayed photosynthetic induction in *G. max*, which was achieved by developing a cold-girdling system using the Peltier device that can induce NSCs accumulation with minimum effects on N supply and the whole-plant growth. The transcriptome analysis revealed that NSCs-induced photosynthetic downregulation was explained by coordinated downregulation of genes involved in photosynthetic processes, stomatal opening, and nitrogen assimilation in *G. max*. We further investigated how sink-source imbalance causes accumulation of sugars and starch in source leaves, suggesting that downregulation of *TPT* and upregulation of *GPT*s in chloroplast membranes promote the chloroplast-directed transport of hexose phosphate, which promote the starch accumulation in the chloroplast. It is being predicted that photosynthetic downregulation due to NSCs accumulation will become more pronounced in the future high CO_2_ environments, which will have a serious impact on crop production. Therefore, based on the present results, it is expected to create/breed crops with increased sink activity and sugar translocation capacity as well as enhanced nitrogen absorption and assimilation capacity to cope with elevated CO_2_.

## Materials and methods

### Plant material and growth conditions

Seeds of soybean (*Glycine max* (L.) Merr. ‘Tsurunoko’) and common bean (*Phaseolus vulgaris* L. ‘Yamashiro-kurosando’) were purchased from a commercial supplier (Takii seed, Kyoto, Japan). Seeds were placed on wet paper and kept at 4°C overnight and then transferred to a growth chamber (FHC-740, Espec Mic Corporation, Osaka, Japan). Plants were grown at a photosynthetically active photon flux density (PPFD) of 200 μmol m^−2^ s^−1^ provided by LED during 12-h light period. Temperatures and relative humidity were set to 25°C and 60%, respectively, and CO_2_ concentrations was controlled at 400 μmol mol^−1^. Germinated seedlings were transplanted in 300-mL plastic pots filled with vermiculite at 5 to 7 days after transferring to the growth chamber. During the growth period, pots were randomly rotated every two or three days.

The nutrient solution contained 1 mM NaH_2_PO_4_, 0.25 mM Na_2_HPO_4_, 1 mM MgSO_4_, 10 μM Fe-EDTA, 0.1 mM MnSO_4_, 0.3 mM H_3_BO_3_, 10 μM ZnSO_4_, 1 μM CuSO_4_, 0.25 μM (NH_4_)_6_Mo_7_O_24_, and 1.25 μM CoCl_2_. High (10 mM N) and low nitrogen (0 mM) solutions with 2 mM K and Ca concentrations were obtained by adding KNO_3_, Ca (NO_3_)_2_, KCl, and CaCl_2_. After transplanting, plants were irrigated with 50 ml of high N solution every day for about 14 days. until the primary leaves fully expanded.

### Experimental design

The experimental design was described in Figure S1. The day when primary leaves fully expanded was defined as 0 days after treatment (DAT), and the first photosynthesis measurements were conducted as described below. From 0 DAT, plants were divided into four groups: Control (Ct), cold-girdling (CG), sucrose-feeding (Suc), and low nitrogen treatments (LN), which were intended to accumulate NSCs in primary leaves. In CG treatment, a 3 cm × 3 cm aluminium plate with Peltier element (Denshi Tsusho Corporation, Tokyo, Japan) was attached to the main stem below the primary leaves, and surface temperature of the device was controlled at 3-5°C to suppress phloem transport. In Suc treatment, 50 ml of 20 mM sucrose solution was applied to the plants in addition to high N solution every day. In LN treatment, 50 ml of low N solution was applied to the plants every day instead of high N solution. Plants in the Ct, CG and Suc treatments were irrigated with 50 ml of high N solution from 1 DAT to 3 DAT and 100 ml per pot from 4 DAT to 7-8 DAT.

We conducted the experiments three times. We examined changes in photosynthetic characteristics in response to NSCs accumulation caused by CG treatment (Exp. 1) and LN and Suc treatments (Exp. 2) in *G. max* and *P. vulgaris*. In Exp. 3, we further investigated transcriptomic changes in response to NSCs accumulation caused by CG and LN treatments in *G. max*.

### Leaf gas exchange measurement

Temporal response of net CO_2_ assimilation (*A*) and stomatal conductance (*g*_s_) to a step change in PPFD from 200 to 1000 μmol m^−2^ s^−1^ in primary leaves were determined using a portable photosynthesis measurement system (LI-6800; Li-Cor BioSciences, Lincoln, NE, USA). In Exp. 1, the measurement was conducted at 0, 2, 4, and 5 DAT for *G. max* and 0, 3, 5, and 7 DAT for *P. vulgaris*; in Exp. 2, at 0, 2, 4, 5, and 7 DAT for *G. max* and 0, 3, 6, and 7 DAT for *P. vulgaris*; and in Exp. 3, 0, 7 DAT or 8 DAT for *G. max*. The measurement started just after plants were transferred from the growth chamber to a measurement room. The leaf chamber was maintained at 400 μmol mol^−1^ CO_2_ concentration (C_a_), a leaf temperature of 25°C, and RH of 50%. The primary leaves were irradiated at PPFD of 200 μmol m^−2^ s^−1^ for 5 min, which was almost equal to the growth light intensity, then PPFD was increased to 1000 μmol m^−2^ s^−1^ for 25 min. The maximum photosynthetic rate (*A*_max_), the maximum stomatal conductance (*g*_smax_), and the time required to reach 90% of *A*_max_ and *g*_smax_ defined as *T*_90A_ and *T*_90*g*s_, respectively, during the photosynthetic induction were determined to characterise the photosynthetic traits (Fig. S2).

### Sampling

Plants were harvested in the evening after the photosynthesis measurements on the last day of each experiment. Three leaf discs (1 cm in diameter) were sampled from the primary leaves used for the gas exchange measurements, oven-dried at 80°C with the rest of the leaves, and finely ground using a mill (TissueLyser II, Retsch, Haan, Germany) for later determination of LMA (leaf mass per area), leaf nitrogen content, and NSCs in Exp. 1, 2, and 3. Another primary leaf not used for the photosynthesis measurements were immediately frozen in liquid nitrogen and stored at –80°C for later transcriptome analysis in Exp. 3. Plant was further divided into the rest of the leaves (upper leaves), stems, and roots. They were oven-dried at 80°C, weighed to determine dry mass and leaf-to-root ratio (leaf dry mass per root dry mass), and finely ground using the TissueLyser II for later determination of nitrogen content and NSCs. Nitrogen content (N_mass_, g N g^−1^) in the leaves, stems, and roots were determined with a CN analyzer (Vario EL III, Elementar, Hanau, Germany), and leaf nitrogen content per area (N_area_, g N m^−2^) was calculated as a product of N_mass_ and LMA. Whole-plant N_mass_ was calculated as whole-plant nitrogen content divided by whole-plant biomass.

### Determination of NSCs

The ground samples (3–5 mg each) were used to determine the contents of glucose, sucrose, and starch as in Araya et al. (2006). Soluble sugars were extracted with 80% ethanol, and sucrose was hydrolyzed to glucose and fructose with an invertase solution (Wako Chemical). The precipitate was treated with amyloglucosidase (cat. no. A-9228; SigmaAldrich) to break down starch into glucose. Finally, glucose, and glucose equivalents of sucrose and starch, were quantified using a Glucose CII Test Kit (Wako Chemicals) and a micro plate reader (Sunrise Rainbow Thermo, Tecan, Männedorf, Switzerland). Whole-plant NSC_mass_ was calculated as whole-plant NSCs content divided by whole-plant biomass.

### RNA-seq analysis and expression analysis of key genes

The frozen primary leaves obtained in Exp. 3 of *G. max* were ground with TissueLyser II. Total RNA was extracted using RNeasy Plant Mini Kit (QIAGEN, Hilden, Germany) and purified using DNase. After checking RNA concentration with a Qubit 2.0 Fluorometer (Thermo Fisher Scientific, MA, USA), cDNA libraries were constructed from 1 μg/sample of total RNA using library preparation kits (NEBNext Ultra II RNA Library Prep Kit for Illumina 7770L, NEBNext Multiplex Oligos for Illumina E7335S, NEBNext Poly(A) mRNA Magnetic Isolation Module E7490L, New England Biolabs, MA, USA) and purified with magnetic beads (Oligo d(T)_25_ Magnetic Beads, New England Biolabs, MA, USA; AMPure XP, BECKMAN COULTER, CA, USA) following the manufacturer’s instructions.

These cDNA libraries were sequenced using NextSeq 500 (Illumina, CA, USA), and the reads were mapped to the soybean reference genome (Wm82.a2.v1). The number of reads mapped to each reference was counted, and the read counts were processed with edgeR and analysed using the iDEP 9.0 (http://bioinformatics.sdstate.edu/idep/), and principal component analysis (PCA), k-means clustering analysis, and pathway enrichment analysis were conducted. Transcript levels of key genes for photosynthesis, stomatal responses, nitrogen assimilation, starch synthesis/degradation, and sugar transport/metabolism were assessed in a detailed manner using the RNA-seq data of *G. max*.

Since *G. max* has many duplicated genes (Shoemaker et al., 2006), we calculated the transcript levels of genes, which are annotated with Arabidopsis gene ID, as the total read counts of duplicated genes based on the assumption that only a small proportion of the duplicated genes obtained new functions or lost their function (Roulin et al., 2013). We focused on the key enzymes in the Calvin cycle responsible for carbon fixation and the factors involved in the activation and assembly of Rubisco. As for stomatal responses, transcript levels of photoreceptors and ion channel/transporter preferentially expressed in guard cells and promoting stomatal opening/closure were investigated. We further examined the expression profiles of nitrate transport, nitrogen assimilation, and nitrogen remobilization and those involved in the sugar-phosphate translocators in the chloroplast envelope.

## Supplemental data

**Supplemental Figure S1**. Experimental design of the present study.

**Supplemental Figure S2**. Theoretical temporal response of net CO_2_ assimilation (*A*) and stomatal conductance (*g*_s_) to a step change in PPFD from 200 (shaded area) to 1000 (unshaded area) μmol m^−2^ s^−1^.

**Supplemental Figure S3**. Photosynthetic traits (*A*_max_ and *g*_smax_) in *G. max* and *P. vulgaris* from 0 to 7–8 days after treatments (DAT).

**Supplemental Figure S4**. Photosynthetic traits, nonstructural carbohydrate (NSC_area_) and leaf nitrogen content per area (N_area_) in the primary leaves of *G. max* in Exp. 3.

**Supplemental Figure S5**. Relationships between photosynthetic traits (*A*_max_ and *g*_smax_), nonstructural carbohydrate (NSC_area_) and leaf nitrogen content per area (N_area_) in the primary leaves of *P*. vulgaris on 7 days after treatments (DAT).

**Supplemental Figure S6**. Principal component analysis (PCA) of transcriptome data of *G. max* in Exp. 3.

**Supplemental Table S1**. Information of genes expressed in each cluster.

**Supplemental Table S2**. Theoretical temporal response of net CO_2_ assimilation (*A*) and stomatal conductance (*g*_s_) to a step change in PPFD from 200 (shaded area) to 1000 (unshaded area) μmol m^−2^ s^−1^.

**Supplemental Table S3**. Photosynthetic traits (*A*_max_ and *g*_smax_) in *G. max* and *P. vulgaris* from 0 to 7–8 days after treatments (DAT).

**Supplemental Table S2. N**ormalized transcript levels of genes in each cluster.

**Supplemental Table S3**. Log2 fold change (FC) in transcript levels of key genes in Calvin cycle.

**Supplemental Table S4**. Log2 fold change (FC) in transcript levels of genes of subunits of photosystems.

**Supplemental Table S5**. Log2 fold change (FC) in transcript levels of genes of light harvesting complexes of photosystems.

**Supplemental Table S6**. Log2 fold change (FC) in transcript levels of genes of cytochrome b_6_f complex.

**Supplemental Table S7**. Log2 fold change (FC) in transcript levels of genes promoting stomatal opening or closure.

**Supplemental Table S8**. Log2 fold change (FC) in transcript levels of genes involved nitrogen transport and assimilation.

**Supplemental Table S9**. Log2 fold change (FC) in transcript levels of genes invloved in starch synthesis in chloroplast and sucrose synthesis in cytosol.

**Supplemental Table S10**. Log2 fold change (FC) in transcript levels of genes invloved in starch degradation.

**Supplemental Table S11**. Log2 fold change (FC) in transcript levels of genes invloved in sugar transport.

**Supplemental Table S12**. Log2 fold change (FC) in transcript levels of genes proposed to be involved in sugar-sensing.

**Supplemental Table S13**. Log2 fold change (FC) in transcript levels of senescence-associated genes.

## Acknowledgement

We thank the members of Crop Science laboratory in Nagoya University, Dr. Ichiro Terashima, and Dr. Mizokami for their valuable comments and encouragement.

## Author contributions

YO and DS designed the experiments. YO cultivated the plants and conducted the gas exchange and physiological measurements. YO, AT, and DS contributed to the preparation of RNA-seq samples, and TS performed the RNA sequencing and data analyses. YO and DS interpreted the data and wrote the manuscript.

